# Effects of siRNA silencing on the susceptibility of the fish cell line CHSE-214 to *Yersinia ruckeri*

**DOI:** 10.1101/626812

**Authors:** Simon Menanteau-Ledouble, Oskar Schachner, Mark L. Lawrence, Mansour El-Matbouli

## Abstract

Bacterial pathogens are known to co-opt mechanisms of the host cells’ physiology to gain intracellular entrance. Among the facultative intracellular bacteria is *Yersinia ruckeri*, an enterobacterium mostly known as the causative agent of enteric redmouth disease in salmonid fish. In the present study, we applied RNA inhibition to silence twenty pre-selected genes on the genome of a fish cell line (CHSE-214) followed by a gentamycin assay to quantify the effect of this silencing on the susceptibility of the cells to infection. It was found that silencing of 16 out of 20 genes significantly reduced the number of *Y. ruckeri* recovered at the end of the gentamycin assay. Among the genes with the strongest impact were Rab1A, actin and Rac1, which is consistent with our previous findings that N-acetylcysteine, a chemical inhibitor of Rac1, completely prevented invasion of cells by *Y. ruckeri*. Conversely, silencing of the Rho GTPase activating protein had no statistically significant effect, possibly because *Y. ruckeri*, like some other members of the *Yersinia* genus is able to activate Rho GTPase directly. Similarly, the effect of silencing E-cadherin was not statistically significant, suggesting that this might not be a target for the adhesion molecules of *Y. ruckeri*. Taken together, these findings improve our understanding of the infection process by *Y. ruckeri* and of the interactions between this bacterial pathogen and host cells.

**Importance:** Intracellular invasiveness is a mean for bacterial pathogen to gain shelter from the immune system as well as access nutrients. The enterobacterium *Y. ruckeri* is well characterised as a facultative intracellular pathogen. However, the mechanisms of invasion scrutiny. Investigations have mostly focused on the bacterial virulence rather than on the host’s mechanisms hicjacked during invasion. The present findings therefore allow us to better understand the interaction between this important potentially zoonotic pathogen of fish and host cells in vitro.

## Introduction

The enterobacterium *Yersinia ruckeri* is a major fish pathogen world-wide; and has mostly been studied has the causative agent of enteric red-mouth disease in salmonid fish (1, 2). It is associated with acute outbreaks, especially in younger fish, and causes septicaemia and haemorrhages leading to high levels of mortality in infected fish (1–3), especially at temperature between 18 and 20°C. The bacterium has zoonotic potential and has been associated with topical infections in humans (4). Like several other enterobacteriaceae including other members of the genus *Yersinia, Y. ruckeri* has demonstrated the ability to survive within macrophages (5) as well as invade non-professional phagocytic cells (6, 7). This likely allows the access to some restricted nutrients and protects it from the immune system, moreover it might play a role in the bacterium’s crossing of epithelial membranes as known from other members of the genus *Yersinia* (8).

Two main mechanisms of entry have been described in bacteria, and both are present in *Enterobacteriaceae*. In the trigger mechanisms, effector proteins secreted through the type three and type four secretion system (T3SS and T4SS) interact with regulatory proteins, in particular members of the Rho family (RhoGTPases Rac, Cdc42 or RhoG) (9). This leads to a rearrangement of the cytoskeleton of the host cells resulting in the uptake of the bacterium (10, 11). Interestingly, while *Y. pestis* is known to harbour a T3SS and some of its effector proteins have been shown to target the cytoskeleton and actin filaments. However, these effectors seem to play a role in preventing phagocytosis by professional phagocytic cells rather than to promote intracellular invasion (12). However, the T3SS of *Y. ruckeri* actually belongs to the Ysa family of T3SS, a different family than the one present in *Y. pestis*. It could therefore play a different role in the virulence of *Y. ruckeri*. Indeed, Ysa is more homologous to the T3SS carried on the *Salmonella* pathogenicity island 1 (SPI-1) of *Salmonella enterica* (13, 14). Among the proteins carried on the SPI-1 is the chaperon Invasion protein B (InvB) and the SPI-1 is known to play a role in the intracellular invasion of *S. enterica* (15, 16) so the Ysa of *Y. ruckeri* could plausibly be involved in intracellular invasion of *Y. ruckeri.* However, our knowledge of the *Ysa* T3SS, and that of *Y. ruckeri* in particular is still incomplete (17) and no conclusion is currently possible.

The other most studied mechanism of entry is the zipper mechanism that is considered as the main mechanism of entry for bacteria belonging to the genus *Yersinia.* It is initiated by the binding of the bacteria to specific molecules on the cell membrane. For example, in *Y. pestis*, the attachment-invasion locus (ail) of *Yersinia pestis* is a 17.5 kDa outer membrane protein that plays multiple roles in virulence (18), including a minor role in binding to fibronectin (12, 19, 20). Similarly, the Plasminogen activator (Pla) has been shown to have adhesive property for laminin (21) and likely moonlights as an adhesion factor. In *Y. pseudotuberculosis* and *Y. enterocolitica*, the membrane proteins invasin (Inv) and Yersinia adhesin A (YadA) interact with integrin receptors (22). Interactions of the bacteria with the cell surface receptors lead to the recruitment of more receptors, activation of Rac1 and cytoskeletal rearrangement culminating in the uptake of the bacterium (23).

An important feature of both invasion mechanisms is that they require active uptake of the bacterium by the cell and it is possible to prevent host cells from internalising the bacteria, for example by treating them with chemical blockers (24). This is for example the case for *Y. ruckeri* and several chemical blockers have been shown to prevent bacterial invasion (6, 7).

Beside chemical blockers, silencing of the genes involved with internalisation has also been shown to have inhibitory effects. For example, a screening was performed using 23300 dsRNA to silence over 95% of the annotated genes on the genome of drosophila. This allowed to identify 305 genes whose inhibition interfered with the ability of *Listeria monocytogenes* to invade cell cultures of drosophila SL2 cells and 86 genes necessary for invasion by *Mycobacterium fortuitum* (25, 26). Interestingly, a screening by Kumar *et al.* (27) on 18,174 genes from the genome of human macrophages identified no less than 270 molecules whose silencing affected the load of seven different field isolates of *Mycobacterium tuberculosis* within human macrophages (27). Among these, 74 proteins were involved in the intracellular survival of every isolate tested and over half of the genes identified played a role in the regulation of autophagy, suggesting that this process was central in controlling intracellular *M. tuberculosis*. Conversely, when Thornbrough *et al*. applied a gentamicin assay to investigate the effect of a siRNA library targeting approximately 22.000 genes, they found that a larger variety of genes were involved in the intracellular survival of *Salmonella enterica* serovar *Typhimurium* within the human epithelial cell line MCF-7 (28). Among the 252 genes whose silencing reduced the survival of *S. enterica Typhimurium* more significantly were SEC22A, Rab1B and VPS33B that were involved in vesicle trafficking as well as ATP6VOD1, an ATPase involved in vacuole acidification and the iron transporter FTHL17. Nonetheless, the authors noted a significant overlap between their findings and that reported by Kumar *et al*. for *M. tuberculosis* (27) suggesting that many of these targets are well conserved even between very evolutionary distant bacteria.

In the present study, we aimed to produce siRNA to silence 20 genes selected because they are commonly associated with the intracellular survival of bacteria. The fish cell line CHSE-214 was used and the effect of this silencing on the susceptibility of the cells to intracellular invasion by the facultative intracellular fish pathogen *Y. ruckeri* was investigated using a gentamicin assay.

## Results and Discussion

### Several notable genes were found to be necessary for the intracellular invasion process

Silencing of every one of these genes resulted in reduction of the number of bacteria recovered from Chinook Salmon Embryo (CHSE-214) cells at the end of the assay; however, this reduction was only statistically significant in 16 out of 20 genes (Table 1).

**Tables 1.** Gene investigated in this study and effect of the silencing on the number of bacteria recovered at the end of the gentamycin assay. Greyed out cells represent genes whose silencing did not significantly impact the gentamycin assay.

| Gene name | Function | Significance |
| --- | --- | --- |
| **Protein kinase C** | Signal transduction | P = 0.000 |
| **Rab-1A** | Vesicular trafficking | P = 0.000 |
| **SEC22b-B** | Vesicular trafficking | P = 0.002 |
| **Vacuolar ATP synthase subunit A** | Vacuolar protein sorting | P = 0.003 |
| **VPS-associated protein 33A** | Vacuolar acidification | P = 0.002 |
| **Rho1-GTPase** | Cytoskeleton organisation | P = 0.006 |
| **Ubiquitin conjugating enzyme E2L3** | Tagging of protein for ubiquitin degradation | P = 0.003 |
| **Sumo 2** | Post-translational modification system | P = 0.005 |
| **Equilibrative nucleoside transporter 1** | Cellular uptake of nucleosides | P = 0.000 |
| **Integrin β-1 precursor** | Integrin | P = 0.000 |
| **Actin** | Cytoskeletal apparatus | P = 0.000 |
| **Rac1** | Cell growth & cytoskeletal reorganization | P = 0.000 |
| **SDC42** | Rearrangement of actin filaments | P = 0.001 |
| **Rho GTPase-activating prot.** | Rearrangement of actin filaments | P = 0.795 |
| **Laminin** | Structural scaffolding of the tissues | P = 0.006 |
| **E-cadherin** | Cell adhesion | P = 0.722 |
| **Myotubularin-related protein 2** | Vesicular traffic and actin remodeling | P = 0.045 |
| ***p38b1* mitogen activated protein kinase** | Stress resistance and apoptosis | P = 0.000 |
| **Caspase I precursor** | Role in apoptosis | P = 0.962 |
| **Cyclin D1** | Coordination of mitosis | P = 0.948 |

Among the genes whose silencing had the strongest effect were the protein kinase C, a regulatory protein overseeing a large number of varied functions, including signal transduction as well as the expression of the cytoplasmic tyrosine kinase, Focal Adhesion Kinase (FAK) and actin rearrangement (29). Previously, protein kinase C activity has shown an inverse correlation with the uptake of *L. monocytogenes* by J774, a cell line of murine macrophage, as well as a positive correlation with the escape of these bacteria from the macrophages endosomes (30). Enteropathogenic *Escherichia coli* is also known to recruit protein kinase C to the membrane of HeLa as well as the colon carcinoma cell line T84 (31) and more recently, it has been shown that protein kinase C recruitment in lipid rafts was induced by Enterohemorrhagic *E. coli* O157:H7. In the present study, silencing of the gene encoding for protein kinase C resulted in a strong decrease of the cell susceptibility to bacterial infection (P=0.000).

Another important gene was the Ras related protein Rab1A. The Rab family constitute a subset of the Ras superfamily of proteins which are small GTPase acting together as network involved in the regulation of vesicular transport (32). In humans, more than 60 members of the Rab family have been identified (32) and about 52 in the channel catfish (*Ictalurus punctatus*), although no census of Rab proteins has yet been conducted for salmonid fish (33). Rab1A specifically is required for the microtubule-based motility of murine endocytic vesicles (34) and silencing of Rab1b was shown to lead to a reduced growth of *S. enterica Typhimurium* within MCF-7 cells (28).

Equilibrative nucleoside transporter 1 is a member of a well conserved family of transmembrane proteins and silencing of the corresponding gene was found in the present study to significantly decrease intracellular bacterial survival and multiplication (P = 0.000). Pathogenic intracellular bacteria are known to scavenge nucleotide from their environment and therefore, this difference could be explained by a reduction in the nutrients available to the bacterium (35).

Another molecule whose silencing had a significant effect oncell sensitivity to infection (P = 0.000) was the precursor for integrin β-1. Integrins are trans-membrane receptors and their role in cell adhesion to the extra-cellular matrix is well known. Moreover, integrins β-1 have been well studied as target of the adhesin molecules of *Yersinia* sp., including *Y. pseudotuberculosis* (36) and *Y. enterocolitica* (37). As such, they play a central role in the use of the zipper mechanisms by these bacteria to gain intracellular entry. The finding that this target is also important for *Y. ruckeri* suggests that this bacterium might gain intracellular entry through the zipper mechanism targeting of integrin β, in a manner reminiscent of that of *Y. pseudotuberculosis* and *Y. enterocolitica*.

Similarly, silencing of the actin gene also had a very notable effect (P = 0.000). Because actin is such a central component of the cytoskeleton and because of the critical role of the cytoskeleton in both the trigger and zipper mechanism (11), this finding was not unexpected.

Finally, another molecule of importance was Ras-related C3 botulinum toxin substrate 1 (Rac1). It is a member of the Rho family of GTPases and plays a central role regulating cytoskeleton rearrangement. It has previously been involved in intracellular invasion of the enterobacteriaceae *Salmonella typhimurium* (38, 39) and is targeted by the effector protein IpgB1 of *Shigella* (40). Among *Yersinia* spp., the picture is more complex as Rac1 is inactivated by the T3SS effector protein YopE expressed among others by *Y. pseudotuberculosis* (41) and *Y. enterocolitica* (42). Notably, this Rac1 suppressive activity is most likely to hinder phagocytosis mediated killing (43) while intracellular invasion with the zipper pathway relies on the CDC42-independent activation of Rac1 (44). Moreover; in a previous experiment, we investigated the effect of multiple chemical and found out that the inhibitor of Rac1 N-acetylcysteine completely inhibited entrance of *Y. ruckeri* into CHSE-214 cells (6).

### Conversely, several genes were identified for which silencing did not result in a statistically significant decrease of the cells’ sensitivity to infection

Among these genes were the Rho GTPase-activating protein (P = 0.795). Rho GTPases act as regulator and facilitator for the activity of Rho-GTPase molecules such as Rac1 and CDC42. Several virulence factors of enterobacteriaceae are known to activate Rho GTPase enzymes directly (45), bypassing RhoGTPase activating proteins in a manner consistent with the present findings. An example of such a protein is the cytolytic necrotising factor (CNFy) of *Y. pseudotuberculosis* (46). Interestingly, a segment of the *Y. ruckeri’s* genome is homologous to a section of the gene encoding for *Y. pseudotuberculosis’* CNFy but it is not currently known if this molecule is active in *Y. ruckeri*.

Another protein whose silencing did not result in a significant reduction of sensitivity to infection was E-cadherin (P = 0.722). E-cadherin acts as the receptor for Internalin used by *Listeria. monocytogenes* (47). However, members of the *Yersinia* genus tend to use different surface receptors such as integrins (22). If E-cadherin is not a target of *Y. ruckeri*’s adhesion molecules, then it would be expected that silencing of the corresponding gene would not result in any significant change susceptibility to intracellular invasion.

The next gene whose silencing had no significant effect on the bacterial invasion was the caspase I precursor (P = 0.962). Caspases take part in the cellular immune response and play an important role in regulating apoptosis (48). Caspase I is known to be inhibited by the YopJ effector molecule of *Y. pestis* (49).

Finally, the last gene whose silencing had no statistically significant effect was the cyclin D1. Cyclin D1 regulates cellular proliferation (50) and its expression is known to be affected by intracellular invasion of *Streptococcus pyogenes* (51). However, there is no evidence suggesting it might play a role in the intracellular invasion of Gram-negative bacteria. Moreover, cyclin D1 is most expressed during the G1 phase of the cell life cycle (50).As our cells were confluent at the time of the intracellular invasion procedure, it is expected that the expression of cyclin D1 was lowered even in the absence of RNA inhibition.

To summarise, we used RNAi to silence 20 selected genes on the genome of CHSE-214 cells. Silencing of 15 of these genes resulted in a significantly reduced susceptibility to invasion of *Y. ruckeri*. The results of this study contribute to our understanding of the invasion mechanisms in this important fish pathogen.

## Materials and methods

### Gene selection and design of the siRNA sequences

Twenty genes were selected based on literature and the fact that homologs of these genes have been implicated in the intracellular invasion of other bacterial pathogens. The selection included genes commonly involved in invasion by bacterial pathogens, including surface integrins as such molecules are often targeted by the invasin molecules expressed by other Yersiniaceae (52–54). Other genes were cytoskeletal molecules and genes involved in vacuolar trafficking and maturation (27, 28) as well as agents of the cytoskeletal apparatus as this plays a central part in the internalization of bacteria through both the zipper and trigger mechanisms (55). The selection also included genes and pathways shared by *L. monocytogenes, M. tuberculosis* as well as *S. typhimurium* (25–28). As these pathways were used by such diverse facultative intracellular pathogens, they might also be involved in the intracellular invasion of *Y. ruckeri*. After selection of the genes, sequences for the siRNAs were designed by Ambion Life Technology using the Silencer Select siRNA design algorithm. For the genes for which no *Oncorhynchus tshawytscha* sequences were available, sequences from other members of the *Oncorhynchus* genomes were used instead. Upon reception, the siRNAs were resuspended to 20 µM and stored at −20°C until use, according to the manufacturer’s instruction.

### siRNA transfection was performed based on the manufacturer’s guidelines

This study made use of CHSE-214 cells. The identity of this cell line derived from Chinook Salmon Embryo has been confirmed by PCR and sequencing and their suitability for gentamycin assay has been confirmed before (6). CHSE-214 cells were grown in 24 well plates at 20°C supplied with 1250 µl Minimum Essential Medium (MEM-glutaMAX™, Gibco) containing 2% FBS, where they reached about 80% confluence one day after seeding.

On the day of the assay, 8 µl of the siRNA solution was added to 400 µl of Opti-MEM medium (Gibco). At the same time, 72 µl of RNAiMax lipofectamine reagent (Sigma Aldrich) was diluted in 1200 µl of Opti-MEM medium. The two solutions were mixed together and incubated 5 minutes at room temperature. Afterwards, the culture medium on top of two adjacent rows of the 24 wells (either rows 3 and 4 or 5 and 6; 8 wells in total) was replaced with 50 µl of the solution and, after 20 minutes, 450 µl of fresh MEM-Glutamax medium was added.

In each plate, as negative control, cells from the 8 wells in column 1 and 2 were processed in the same manner but without the siRNA, serving.

The cells were then incubated at 20°C and due to the temperature lower than other cell lines commonly used for transfection, incubation time was extended to three days.

### Bacterial invasion assay

The gentamycin assay was performed as previously described (6). Briefly, *Y. ruckeri* ATCC 29473 was cultivated overnight in brain heart infusion (BHI, Oxoid). Optical density of the culture was assessed by spectrophotometry and adjusted to an optical density of 0.5 at 600 nm. The bacteria were pelleted down by centrifugation at 3220 g and 20°C (in an Eppendorf Centrifuge 5810 R) before being resuspended in 10 times the original volume of MEM-Glutamax. Then, the culture medium in the wells 1A-C, 3 A-C and 5 A-C was replaced by this bacterial solution (for clarity, the plate loading plan is illustrated on Fig. 1) and the bacteria were left to interact with the cells for five hours at room temperature. In the wells 1D, 3D and 5D, the medium was replaced by fresh MEM-glutamax without bacteria. After that time, the medium was removed, the cells were washed three times with PBS and a fresh volume of MEM-glutamax, supplemented with the antibiotic gentamycin (Sigma-Aldrich) at a concentration of 100 μg.ml^−1^ was added to the wells. The antibiotic was left to act for four hours. Afterwards the cells were washed twice with PBS before replacing the medium supplemented with 1% Triton-X (Sigma-Aldrich). After 10 minutes of exposure to the detergent, the cells were triturated with a micropipette and serially diluted from 10^−1^ to 10^−4^ before being plated onto Brain heart infusion agar (BHIA, Oxoid). Each of the three wells represented a biological replicate and each dilution was plated in technical quadriplicates, meaning that 48 plates were inoculated for each siRNA. The agar plates were incubated at 22°C until clear colonies were visible and counted (generally after 48 hours). The corresponding average CFU per ml value was then calculated. To minimise the effect of the variation between cell and bacterial culture, these values were always compared to that of the control CHSE-214 cells from the same plate (Fig 1).

**Fig 1.**
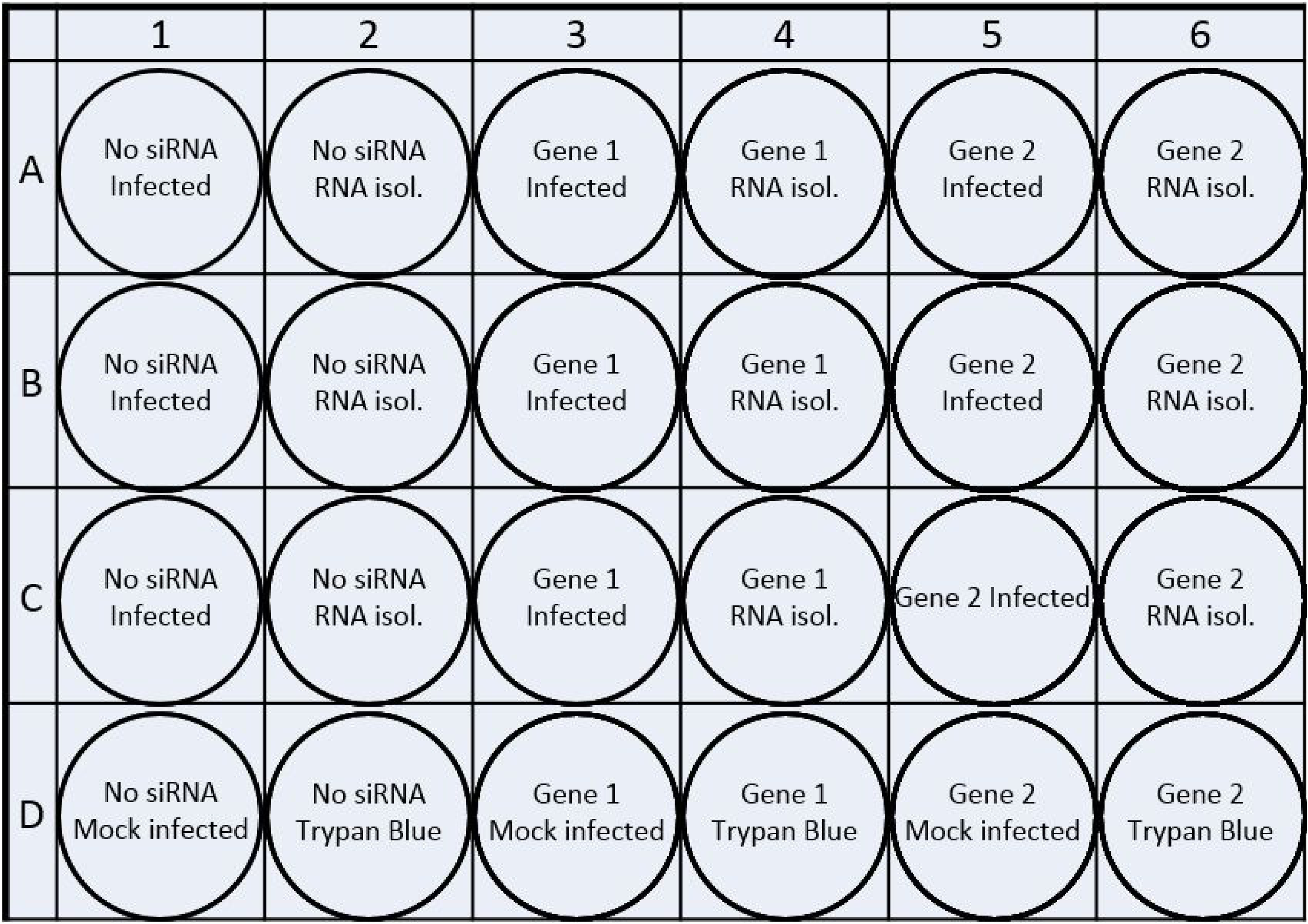
Loading plan for the 24 wells plates used in the gentamycin assay. Each siRNA was tested alongside a non-silenced control to minimise the variations between plates. For each gene silenced, the three first wells of the leftmost row were infected for the gentamycin assay while the bottom well was left uninfected as a control. On the rightmost row, RNA was extracted from the first three wells to confirm silencing by RTqPCR while the last well was stained using trypan blue to assess any toxic effect of the RNA silencing.

### Controls

For each siRNA, in the 24 well plate the cells of number 1D, 3D and 5D were left un-inoculated then lysed and plated to act as a negative control and detect any contamination of the reagents while the cells in 2D, 4D and 6D were exposed tested using a trypan blue assay. Briefly, the cells were washed three times in PBS, then 0.2% Trypan Blue (0.4% diluted 1:1 in PBS) was added. The cells were stained for 1 minutes prior to fixation with 4% formalin for 10 minutes Afterwards they were rinsed several times until any trace of blue dye had disappeared. The plates were kept at 4°C until quantification of the cells using an inverted microscope (Leica DM IRB). One hundred cells were counted and the number of blue stained cells among them was recorded. The procedure was repeated 4 more times for each culture unit to result in 5 percentage values for each siRNA. These were then compared to the survival of the control cells without siRNA to confirm that the silencing procedure did not have a toxic effect. In no instances did these numbers differ significantly from that of the control.

Finally, the cells in the wells 2A-C, 4A-C and 5A-C were lysed in buffer RLT (Qiagen). The cell solution/suspension? was homogenised using QIAshredder columns (Qiagen) and centrifugation at 145000 RPM for two minutes at room temperature using MiniSpin tabletop centrifuge (Eppendorf). Afterwards, the RNA were extracted using the Rneasy mini kit (Qiagen) according to the manufacturer’s handbook. The RNA were immediately quantified using a Nanodrop machine (ThermoFisher) and cDNA were immediately synthesised using Iscript kits (Bio-Rad) according to the manufacturer’s instructions in a C1000 Touch thermocycler (Bio-Rad). Resulting cDNAs were stored at 4°C until use.

### RTqPCRs

The primer sequences used for confirmation of silencing are listed in Table 2. Primers for ubiquitin and elongation factor 1-alpha designed by Peña *et al.* (56) were used for the housekeeping reference genes, as these authors found them among the most suitable when working with CHSE-214.

**Tables 2.**
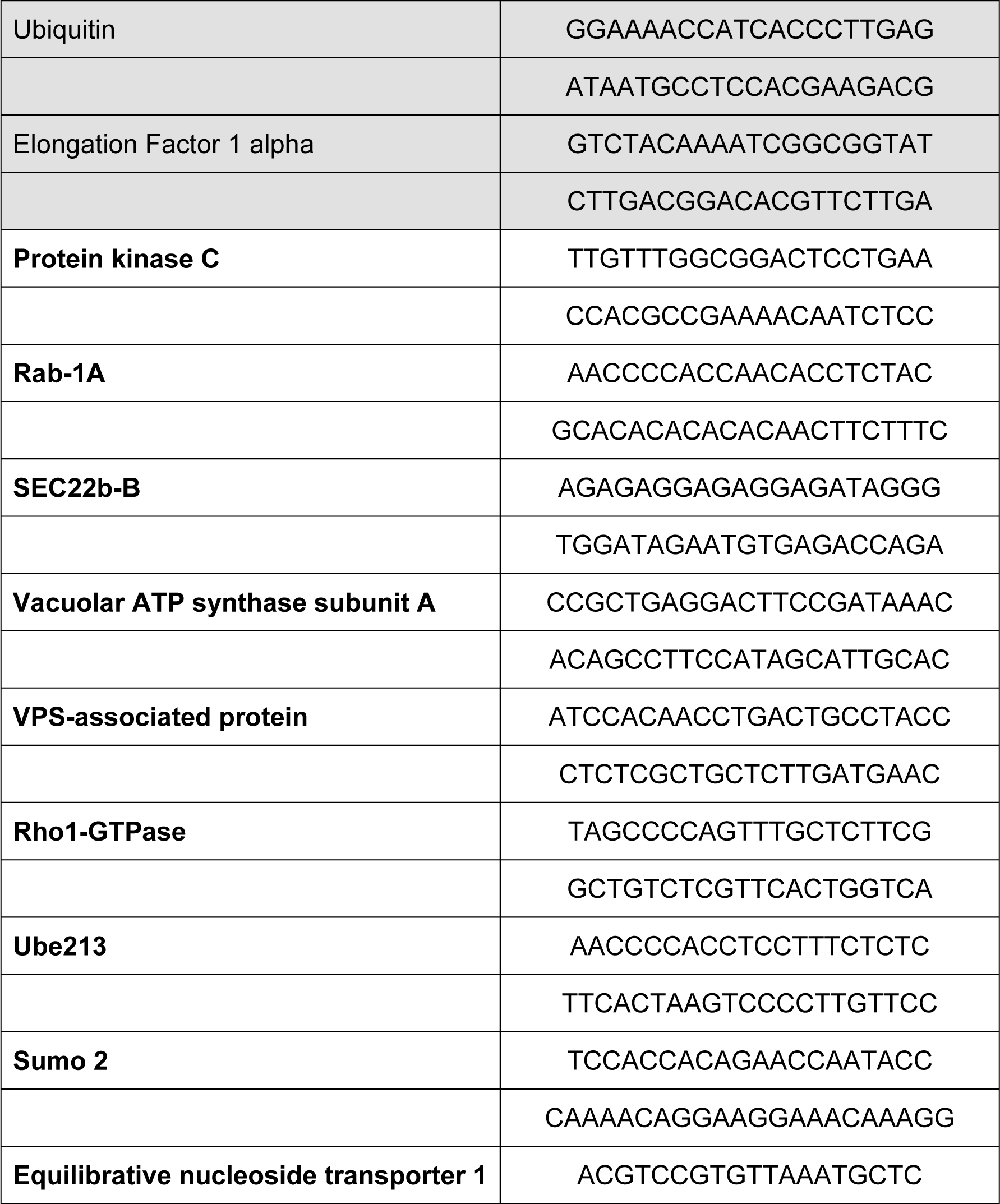
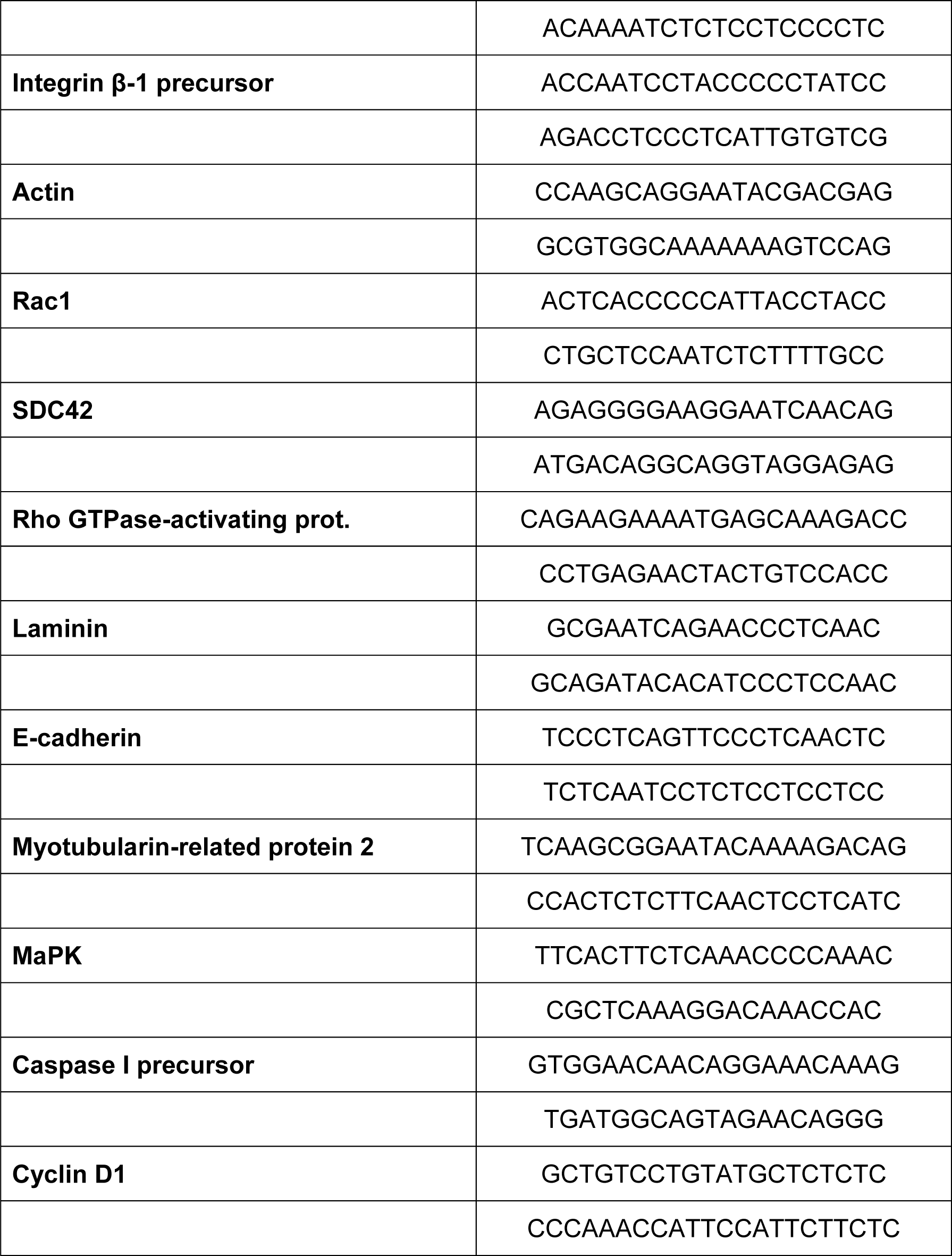
Primer sequences for the RTqPCr confirmation of the siRNA silencing. The two first (greyed out) primer pairs are the housekeeping genes and where designed by Peña *et al.* (56)

Genomic DNA was extracted from CHSE-214 cells using the DNeasy kit (Qiagen) according to the manufacturer’s instructions. End point PCR was performed using these primers to confirm the optimal annealing temperature, afterwards the PCR products were purified with a QIAquick gel purification kit (Qiagen). The products’ concentration were measured using a nanodrop, then adjusted to 10 ng.µl^-1^. Afterwards, serial dilution was performed to produce concentrations ranging from 10^-1^ to 10^-4^ ng.µl^-1^. qPCRs were then performed on these serial dilutions as well as the cDNA produced as described in the previous section. Relative gene expression levels of the silenced genes were calculated using the 2^-ΔΔCt^ method (57) to confirm the efficacy of the silencing. The serial dilutions were used to calculate the R^2^ and efficiency of the qPCRs and confirm that these were above 0.9 and between 95 and 100% respectively for every qPCR.

## Acknowledgments

This research was financed by the Austrian Science Funds (Fonds zur Förderung der wissenschaftlichen Forschung), project P28837-B22. This funding body played no role in the design of the study or interpretation of data.

